# Non-genetic differences underlie variability in proliferation among esophageal epithelial clones

**DOI:** 10.1101/2023.05.31.543080

**Authors:** Raúl A. Reyes Hueros, Rodrigo A. Gier, Sydney M. Shaffer

## Abstract

The growth potential of individual epithelial cells is a key determinant of tissue development, homeostasis, and disease progression. Although it is known that epithelial progenitor cells vary in their proliferative capacity, the cell states underlying these differences are yet to be uncovered. Here we performed clonal tracing through imaging and cellular barcoding of an in vitro model of esophageal epithelial cells (EPC2-hTERT). We found that individual clones possess unique growth and differentiation capacities, with a subset of clones growing exponentially. Further, we discovered that this proliferative potential for a clone is heritable through cell division and can be influenced by extrinsic cues from neighboring cells. Combining barcoding with single-cell RNA-sequencing (scRNA-seq), we identified the cellular states associated with the highly proliferative clones, which include genes in the WNT and PI3K pathways. Importantly, we also identified a subset of cells resembling the highly proliferative cell state in the healthy human esophageal epithelium and, to a greater extent, in esophageal squamous cell carcinoma (ESCC). These findings highlight the physiological relevance of our cell line model, providing insights into the behavior of esophageal epithelial cells during homeostasis and disease.

## Introduction

The stratified esophageal epithelium contains a basal layer of proliferative cells which give rise to increasingly differentiated progeny above. Basal cells must balance proliferation and differentiation to maintain a healthy tissue structure^1–4^. If this balance is disrupted, it can lead to uncontrolled cell division and the emergence of cancer^5,6^. However, the basal cell states underlying the transition between proliferation and differentiation, and how these contribute to differences in clonal growth, remain unknown^2,7–9^.

In the esophageal epithelium, progenitor cells can have different capacities for growth, resulting in cell lineages of different sizes contributing unequally to the tissue^10,11^. In some instances, enhanced growth can be achieved through genetic mutations that alter the equilibrium between proliferation and differentiation, resulting in the generation of a greater number of proliferative progenitors from a single cell^10,12–16^. Thus, clones that acquire these mutations outcompete other neighboring clones that are less proliferative. However, it remains unknown whether the observed variation in growth among epithelial progenitor cells can always be attributed to genetic differences between the cells^8,17,18^.

In other biological systems, populations of cells can have divergent phenotypes even without genetic differences. These differences can stem from differences in gene expression and the accumulation of epigenetic modifications, such as DNA methylation or histone modifications^19–24^. Examples of non-genetic phenotypes are broad and can include therapy resistance^25–29^, cancer cell fitness^30,31^, immune signaling^32–34^, and circadian period^35^. Moreover, in the esophagus, it is also possible for progenitor cells to grow in response to cues in their environment specifically in the setting of injury^7^. Thus, we wondered whether the growth potential of epithelial cells could be mediated by non-genetic differences between cells.

To address this question, we begin by performing a quantitative analysis of the growth dynamics of immortalized human esophagus epithelial cells. When plated at low density, we find that a subpopulation of these cells is highly proliferative, following exponential growth, while the remaining cells show a range of growth potential. Using mathematical modeling, we find that the observed growth dynamics of all clones cannot be explained by a simple two-parameter model of self-renewal and differentiation. While we find that proliferative capacity is not genetic, we use DNA barcoding to show that it is heritable through a limited number of divisions^36^, and by pairing it with scRNA-seq^26,37,38^, we further uncover a distinct transcriptional state present in the highly proliferative clones. Finally, we demonstrate the relevance of these transcriptional states by mapping them to healthy and cancerous human samples. Taken together, our results suggest that a simple progenitor model is inadequate to explain the growth dynamics in this system. Rather, we propose that each clone has a unique proliferative capacity that is heritable through a limited number of cell divisions, and is linked to specific and previously unknown transcriptional states.

## Results

### Individual epithelial clones have unique capacities for growth and differentiation

To study the growth potential of single epithelial progenitor cells, we used an in vitro model consisting of esophageal epithelial cells (EPC2-hTERT) that both proliferate and differentiate in culture. We selected this model to allow us to make quantitative measurements of growth that are not possible in human tissue. To resolve growth from individual cells, we plated the EPC2-hTERT cells at a low clonal density of 50 cells per well in a 6-well plate and allowed them to grow. In this experimental design, the cells are sufficiently spread apart that even when they grow, the clones do not come into contact with each other. We fixed independent wells at discrete time points, including 2, 5, 8, and 11 days, and then quantified the growth of individual clones using imaging.

We considered potential models for how these clones could grow. If all progenitor cells had unlimited growth capacity, they should follow an exponential distribution (until crowding effects dominate the culture). However, given that these cells have the ability to differentiate and then stop proliferating, we expected growth to be governed by two parameters, namely the doubling time of the progenitor cell and the differentiation into the non-proliferative cell. If these two parameters were fixed, we would expect to see every clone reach a similar size and consist of a mixture of proliferative and non-proliferative cells (Fig. 1A, Outcome 1). An alternative model, however, is that each clone has a unique capacity for growth and differentiation, such that the clones grow to a broad range of different sizes (Fig. 1A, Outcome 2).

**Fig. 1.**
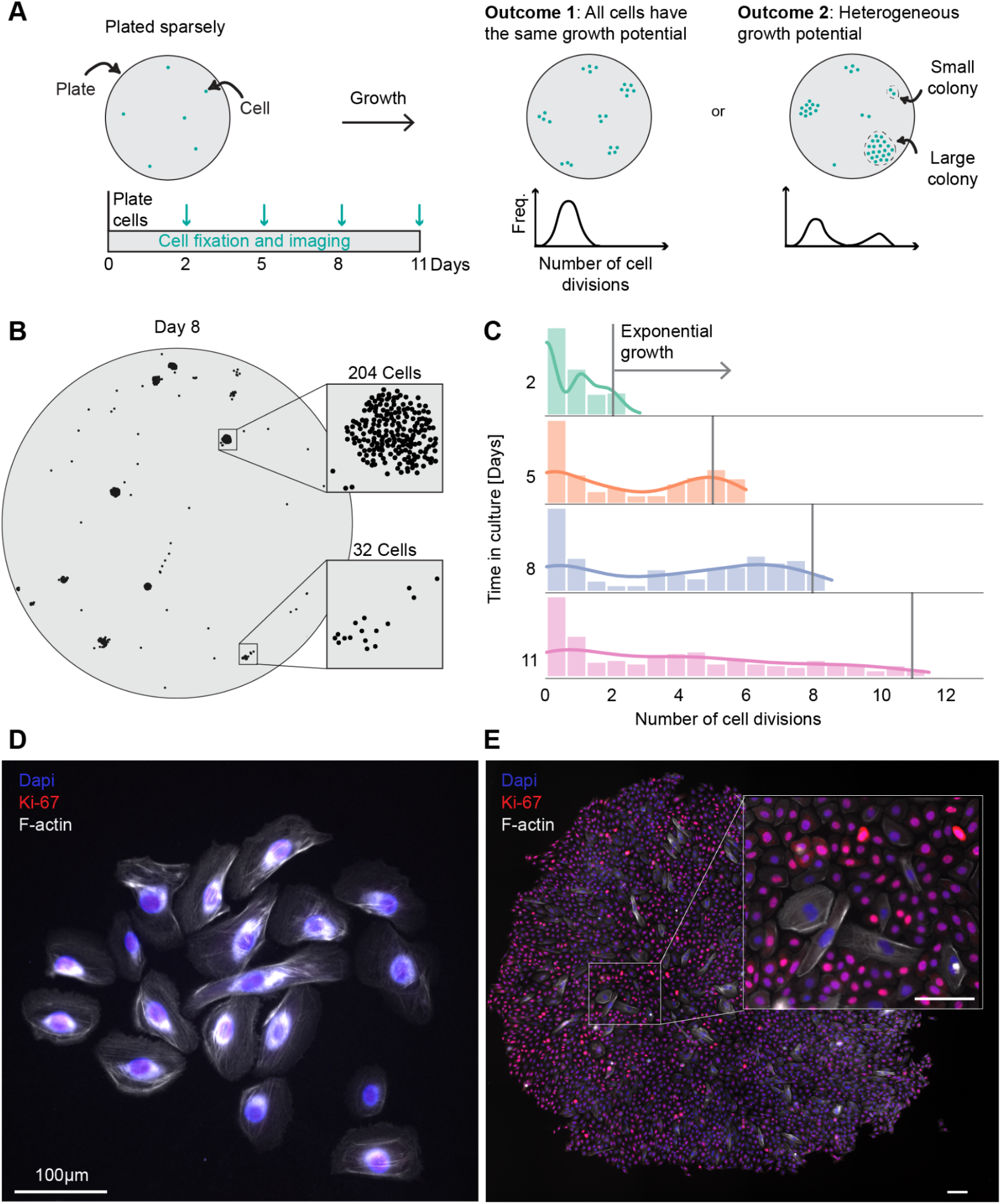
Characterizing cellular proliferation capacity in the esophageal epithelium. (**A**) Schematic of experimental design. On the left, cells are plated at clonal densities, grown for 2, 5, 8 or 11 days and imaged. Possible growth outcomes are on the right. (**B**) A representative plate of sparsely plated cells on day 8. Large and small clones are highlighted on the right with 249 cells and 20 cells, respectively. (**C**) Histogram of the number of cell divisions per clone separated by days in culture. Exponential growth is highlighted with a solid gray line. (**D**,**E**) Immunofluorescence staining for Ki-67 and phalloidin staining for F-actin on day 11 with a small clone on the left and large clone on the right. Scale bars, 100 µm.

Given the sparse plating density, individual clones were easily identified as clusters of cells on the plate (Fig. 1B). By later time points, the clone sizes were highly variable, ranging from 1 to 2,907 cells per clone. To confirm that the larger clones were still growing and the smaller clones had differentiated, we used immunofluorescence to stain for Ki-67, a known proliferation marker, and phalloidin to detect changes in F-actin morphology. We found a distinctive elongation in the cells within the smaller clones; on the other hand, cells forming larger clones, which stained positive for Ki-67, displayed a more rounded and compact morphology (Fig. 1D).

We considered a simple two-state model where a cell can proliferate or transition to a non-proliferative state. We included the non-proliferative state because some clones contain cells that stop proliferating and show morphological features consistent with differentiation. We then performed Gillespie stochastic simulations to generate distributions of clone sizes with a range of parameters (Fig. S1E). We found that this two-state model was unable to fully capture our experimental measurements of clone size within biologically plausible parameter constraints (Fig. S1F). Using a complementary experimental design, we plated single EPC2-hTERT cells in individual wells of a 96-well plate. After 14 days, we found that the majority of clones stopped growing, but approximately 10% of clones continued to grow until they filled the entire well (Fig. S1A, Table S1). Based upon this measurement, we reasoned that 10% of clones grow exponentially, while the others slow or stop growing. Thus, we separated the clone size data (Fig. 1C) into the top 10%, which we named “expanding”. For the remaining clones, we assumed that they followed a pattern consistent with the halting of proliferation, so we named them “committed”. We thus considered the possibility that the highly proliferative clones were a distinct population. The expanding clones grew exponentially, with a reasonable doubling time of 1.732 per day. For the committed clones, a better fit was achieved with a decaying growth rate, since cells transitioned into a non-proliferative state (Fig. S1G,H). Thus, instead of a single two-state model, we show that the growth potential of our esophageal clones is captured by using two independent models, one for each of the proliferative states.

We next considered the possibility that this clonal heterogeneity arose from genetically distinct subclones within the EPC2-hTERTs. To test this hypothesis, we isolated single cells from the EPC2-hTERT line using limiting dilutions in a 96-well plate. We selected the most proliferative clones and then replated them again using a limiting dilution into another 96-well plate. If genetic differences in these clones were responsible for the growth differences, we would expect that all of the subclones derived from the highly proliferative parent clone would grow similarly to the parent. Instead, we found that the subclones followed a similar growth distribution to the initial population (Fig. S1A). Thus, we can conclude that the highly proliferative clones are not a genetically distinct subset of the EPC2-hTERT models.

### Cell states underlying proliferative potential are heritable through cell division

Given that genetic heterogeneity was not the source of these differences in proliferation, we next wondered whether the cell state that underlied the proliferation phenotype was heritable through cell division. To test this hypothesis, we used a high-complexity library of DNA barcodes that are integrated into the genome of single cells and passed on during cell division^38,39^. We transduced the EPC2-hTERT cells with the barcode library at a low MOI to achieve one barcode per cell and allowed them to proliferate through four doublings to generate roughly 16 copies of each barcode in the population. We next replated these barcoded cells randomly into two separate wells at clonal density. We then allowed the sparsely plated barcode clones to grow over 8 days, generating clones of different sizes as we previously observed (Fig. 1B). If the proliferative phenotype were not heritable, we would expect to see clones grow to different sizes in the two wells (Fig. 2A, Outcome 1). On the other hand, if the proliferative potential of a clone were heritable, then the growth of clones across the two wells would be similar (Fig. 2A, Outcome 2).

**Fig. 2.**
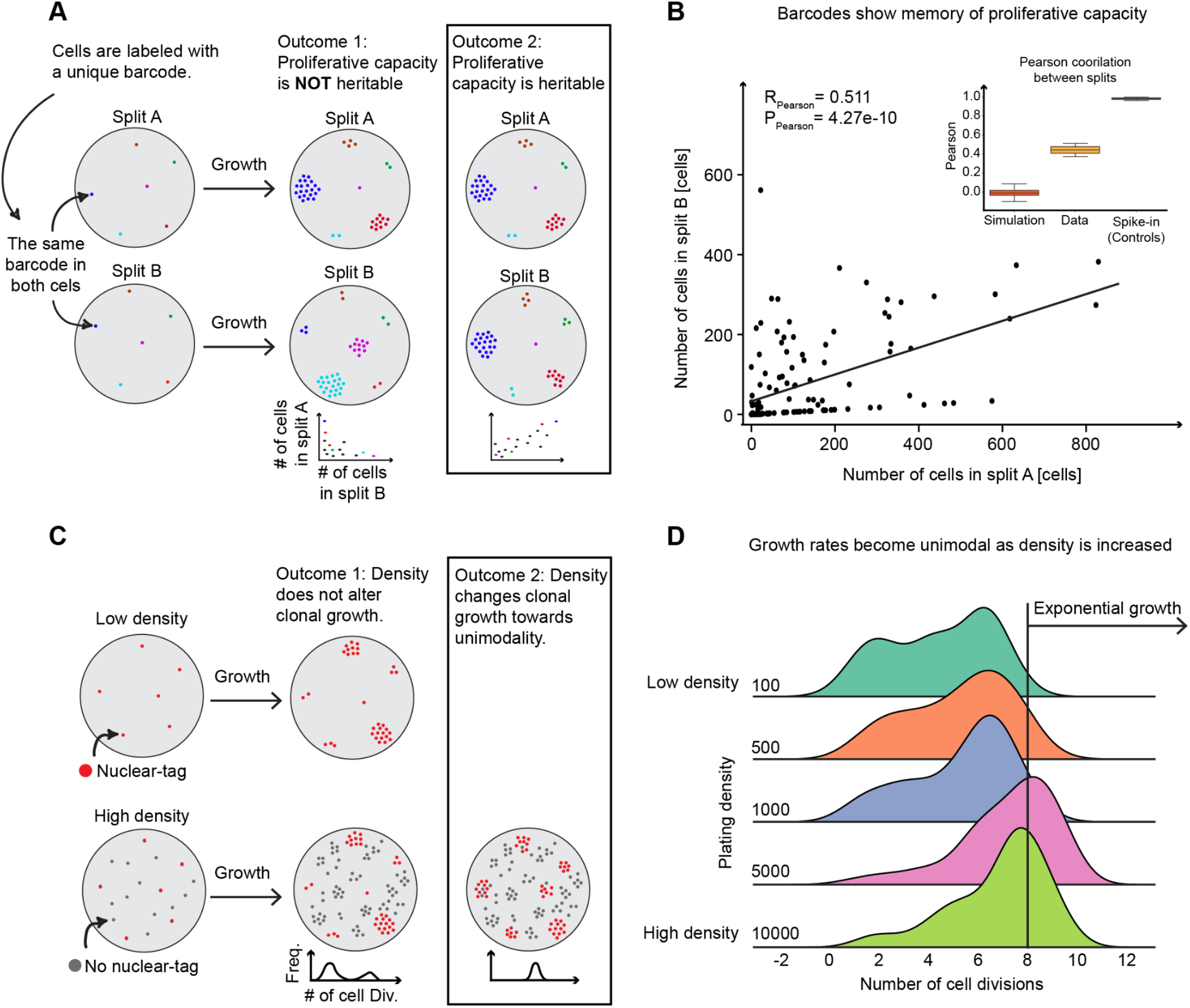
Proliferative capacity is heritable in isolation and can be altered through extrinsic factors. (**A**) Schematic of experimental design showing how lineage barcodes were used to extract information about proliferative capacity. Outcomes of heritable and non-heritable states are described on the right. (**B**) Scatter plot of clonal cells between split A and split B with a Pearson correlation of 0.511 and a p-value of 4.27e-10. The embedded box plots summarize the findings with simulated growth data, three replicates, and spike-in controls. The simulated data was done with bimodal growth rates where each cell picked a random growth rate from the distribution. (**C**) Schematic of experimental design showing mCherry-tagged cells being platted at a constant 50 cells per plate while non-fluorescent cells are seeded at 100, 500, 1000, 5000, or 10000 cells per plate. Outcomes from distinct densities and their effect on proliferation are shown on the right. (**D**) Histograms of cell divisions separated by their original plating density at day 8. Exponential growth is highlighted with a solid gray line.

After eight days of growth, we isolated genomic DNA from both samples to measure the abundance of clones on each plate. We performed a targeted amplification of the barcode sequences and quantified the number of reads for each barcode. When we compared the number of reads in split A and split B, we found a significant correlation in the number of reads for each barcode across the different plates (Fig. 2B). This correlation shows that the proliferative potential of each clone is heritable through cell division, since cells derived from the same parent cell (and labeled by the same barcode) proliferate to similar sizes in independent wells.

To confirm that the number of reads was a good proxy for the number of cells, we added aliquots of defined numbers of cells containing known barcode sequences (50, 500, and 1000) to each sample. We included two samples of each number of cells as replicates. We expect that these replicates should be highly correlated as they were loaded with the same number of cells, and, indeed, they were highly correlated as measured by the Pearson correlation of 0.998 (Fig. S1A). To determine how much correlation we would observe with random growth rates, we simulated the growth of each clone by sampling the experimental growth rates based on their probability density function. The average Pearson correlation with random sampling was 0.012, while our experimental correlation was 0.511 with a p-value of 1.97e-4 between the two. We thus concluded that the proliferative cell state was heritable through the limited number of cell divisions in this experiment.

### Extrinsic cues from neighboring cells can modulate the growth of clones

Having established that heritable cell-intrinsic factors can influence a clone’s growth potential, we next wondered if extrinsic factors could also have an effect. Specifically, we asked whether plating density could alter clone size. However, this posed an experimental challenge because at higher cell densities, it is difficult to accurately track the growth of individual clones. To overcome this challenge, we generated wells with a fixed number of mCherry-labeled EPC2-hTERT cells and varied the number of unlabeled EPC2-hTERTs. In our lowest density condition, we included 50 mCherry labeled cells and 50 unlabeled cells. In our higher density conditions, we maintained the same number of mCherry labeled cells, but varied the unlabeled cells to reach a total of 100, 500, 1000, or 5000 cells per well. We could then track all the labeled cells derived from the same parent cell, as these were located close to each other on the plate. At each density, we allowed the cultures to grow for eight days and then measured the growth of each mCherry-tagged clone by imaging.

If cell density did not alter the propensity for growth, we would expect the size of the clones to be uniform across different densities. However, if higher plating density indeed increased growth, we expected to see increasing clone sizes as density increased (Fig. 2C). Consistent with the previous clonal density experiments, at the lowest density we found a range of different clone sizes, with 10% of clones following an exponential growth curve. As the overall density of the cultures increased, however, we found that the number of cells in each clone also increased (Fig. 2D). We observed the largest average clone size at a plating density of 5,000 cells per well. At the highest density of 10,000 cells per well, the clone size was slightly lower, likely due to the culture approaching over-confluency. Given that most clones became highly proliferative in the higher-density culture, we concluded that the growth potential of individual clones could be modulated by increasing the density of cells, suggesting that external factors in the microenvironment can overcome the intrinsic differences between clones.

### The transcriptional state of expanding and committed cells are distinct

Returning to intrinsic clonal differences, we next wondered whether there were transcriptional differences between the less proliferative and highly proliferative clones. To simultaneously measure clone size and transcriptional state of single cells, we paired DNA barcodes with scRNA-seq (Fig. 3A). We transduced EPC2-hTERT cells with this barcode library, plated them at a low density to capture the non-uniform clonal growth, and allowed them to grow for eight days. Afterward, we collected the cells for scRNA-seq and recovered the transcribed barcodes from the cDNA libraries. By matching the barcodes to the transcriptomes of single cells, we identified clones of different sizes and visualized their locations using uniform manifold approximation and projection (UMAP) (Fig. 3B,C, Fig. S3A,B). We then labeled cells based on their respective clone sizes. Consistent with our previous observations, cells in the top 10% of clone sizes were labeled as expanding, while those in the remaining smaller clones were labeled as committed.

**Fig. 3.**
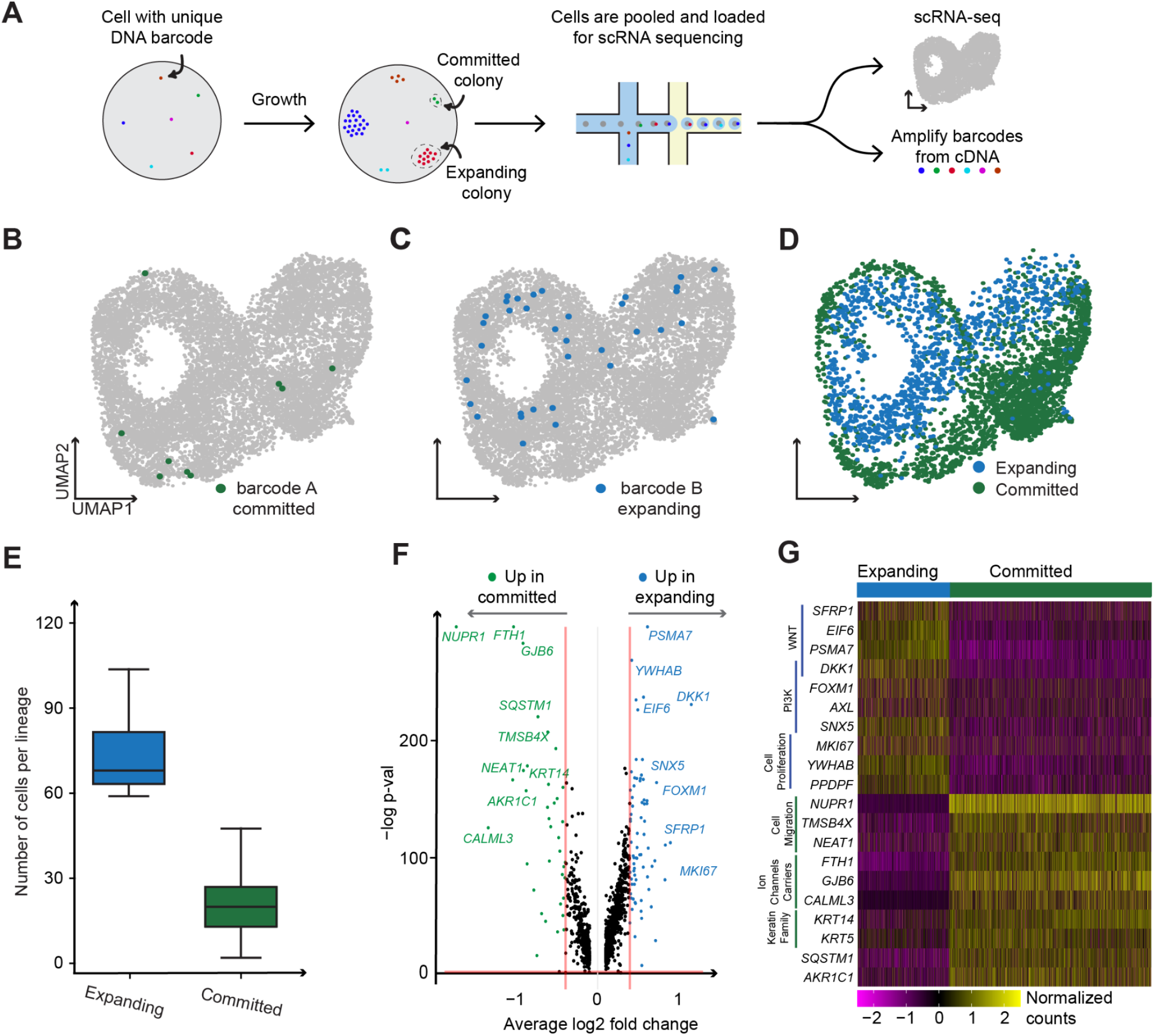
Expanding cells have a unique transcriptional state in which WNT and PI3K pathway genes are upregulated. (**A**) Schematic of how lineage barcodes were used in combination with scRNA-seq. (**B**,**C**) UMAP plots with an example lineage from committed and expanding clones respectively. (**D**) UMAP plots of all lineage barcodes labeled as committed or expanding. Expanding clones make up the top 10% of clones by size, while committed clones were determined by a gene set found through DGE. Unclassified clones didn’t fit either criterion. (**E**) Box plots of the number of cells per lineage in each clone type. (**F**) Volcano plot of differential expression analysis between expanding (positive fold change) and committed (negative fold change). (**G**) Heatmap of the log10 normalized and scaled gene expression of expanding and committed cells.

**Fig. 4.**
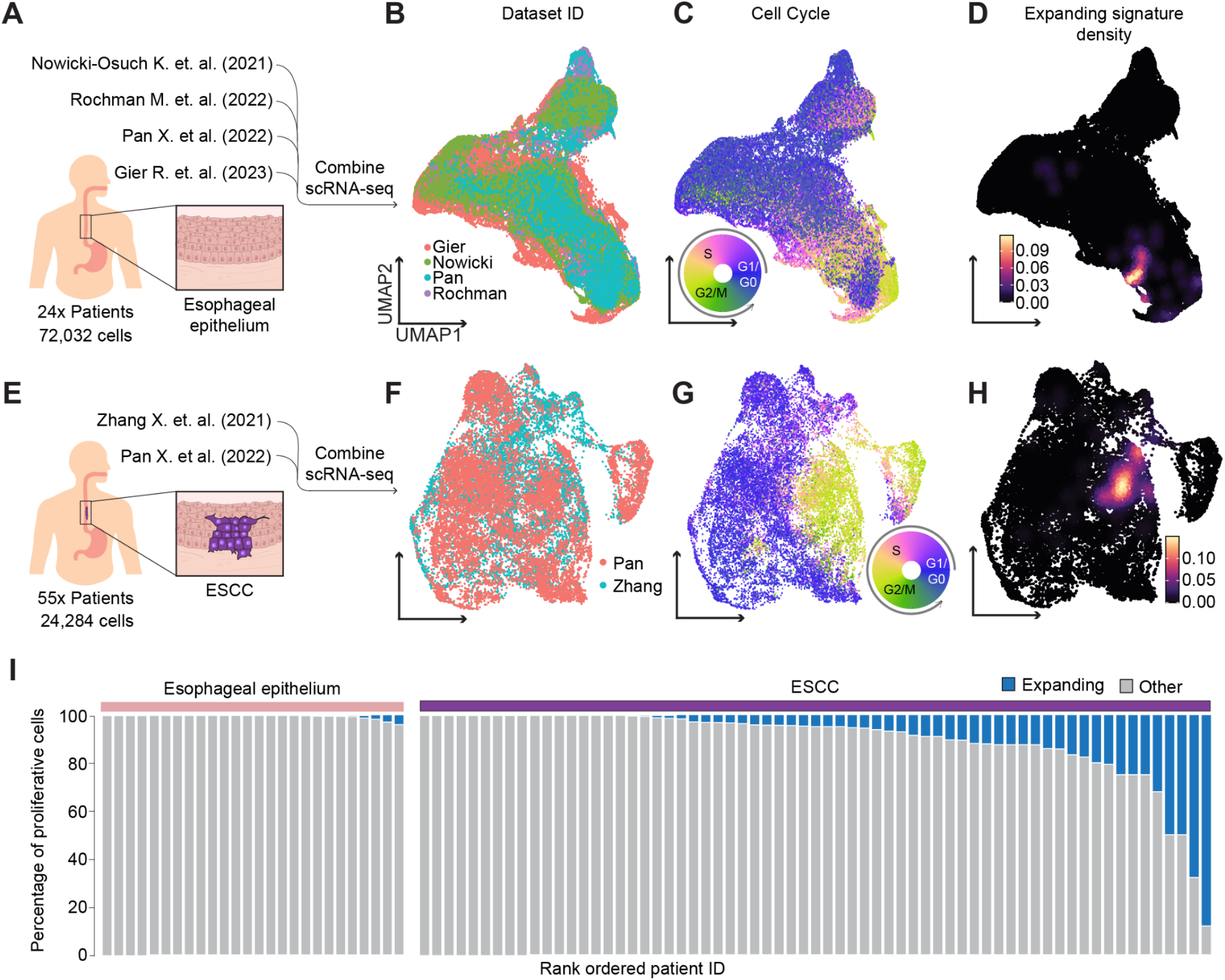
The healthy human esophageal epithelium contains rare basal cells with a similar profile to the expanding clones found in vitro and increase in abundance in ESCC. (**A**,**E**) A schematic of the integration of numerous published scRNA-seq datasets for healthy esophageal epithelium (Top) comprising 26 patients and 62,216 cells, and ESCC (Bottom) with 65 patients and 19,289 cells. (**B**,**F**) UMAP embeddings were corrected for batch effect by patient ID, with clusters colored and labeled according to their original dataset ID. (**C**,**G**) UMAP embeddings, with cells colored based on their cell-cycle position using a circular color scale. Discrete stage labels are placed in approximate positions on the circular legend. (**D**,**H**) UMAP embeddings are overlaid with kernel density estimations showing locations of cells with an expanding signature. (**I**) The stacked colored bars depict the percentage of proliferative cells for each patient, with cells exhibiting an expanding signature in blue and the rest of the proliferative cells in gray. Proliferative cells were selected from their cell-cycle position.

Upon visualizing these expanding and committed populations, we found that they were separated from each other in UMAP space (Fig. 3D,E) and had significant transcriptional differences (Fig 2F). We also noticed that UMAP showed a strong separation of cells based on their predicted phase in the cell cycle (Fig. S3C). However, we found that the cell cycle differences did not align with the differences between expanding and committed clones. Both populations of cells contained cells in each phase of the cell cycle, indicating that the transcriptional differences between these populations are not predominantly driven by cell cycle genes. Of note, we do observe some cells from committed clones that are cycling. This finding is consistent with the decaying growth rate model in which these clones retain some proliferative cells, generated through progenitor cell renewal.

Comparing the transcriptional state of cells in the expanding and committed clones, we found significant differences in gene expression (Fig. 3F, Table S2), with cells from expanding clones having a higher expression of MKI67, as expected. We also found that cells from expanding clones had higher expression of multiple WNT pathway genes, including SFRP1, DKK1, EIF6, and PSMA7, and PI3K pathway genes, including FOXM1, AXL, and SNX5 (Fig. 3G). Intriguingly, a previous study showed that DKK1 and FOXM1 are expressed together in esophageal squamous cell carcinoma (ESCC) and form a positive feedback loop that can promote cell growth^40^. We found this same co-expression pattern in cells from the expanding clones, suggesting that a similar feedback loop could be maintaining the proliferative state in these cells.

While scRNA-seq showed expression of PI3K pathway genes, we next wanted to explicitly test whether the expanding clones had higher PI3K activity and whether this activity was necessary for growth. We used a PI3K FOXO1 reporter which revealed that PI3K activity was robust in expanding clones, whereas committed clones exhibited a decline in PI3K activity (Fig. S4F,G). We next used a PI3K inhibitor, GDC-0941, which resulted in a significant reduction in the total cell count per colony, underscoring the importance of the PI3K pathway in supporting expanding clonal growth (Fig. S4E). To determine whether DKK1 was also important in proliferation, we created a DKK1 knock-out (KO) cell line and found that this knockout increased the percentage of expanding clones (Fig. S4B,C), although it did not make every clone grow exponentially. We thus reasoned that DKK1 could shift the balance between the states regulating the expanding and committed clones. This could be mediated through a redirection of DKK1’s role, possibly by activating non-canonical pathways or mechanisms that promote cell proliferation specifically in expanding clones^40,41^. Together, we find that both PI3K activity and DKK1 can modulate the proliferative phenotypes that underlie these clonal growth behaviors.

### A rare subset of cells with an expanding transcriptional signature was found in the human esophageal epithelium

To determine if the proliferative signatures established in the EPC2-hTERT cell line model was reflected in healthy human esophageal tissue, we collated and analyzed multiple published scRNA-seq datasets of healthy human esophageal epithelium^9,42–44^. After stringent quality control filtering, we combined data from 62,216 cells across 26 patients and batch-corrected the data (Fig. 5A,B). Our primary focus was the basal layer, due to its strong PI3K activity signature that is consistent with our in vitro model. In 4 out of 24 patients, we identified a discernible subset of cells in the basal layer which contained high expression levels of the WNT/PI3K marker genes FOXM1, DKK1, MKI67, and TPX2 (Fig. 5I, left). This subpopulation of cells mirrored the transcriptional profile of expanding clones identified in our cell line.

Shifting our focus to disease states, we turned to ESCC using two scRNA-seq datasets of patients with ESCC^44,45^. Our analysis incorporated data from 19,289 cells across 65 patients (Fig. 5E,F). Compared to the healthy esophagus, ESCC showed a significant increase in the proportion of cells with the expanding clone signature. Across 65 patients total, we found that 33 of the patients had 5% or more of their cells in this transcriptional state (Fig. 5I, right). Taken together, while we find that a small subset of normal patients have cells with this proliferative transcriptional state, this signature is more common in ESCC. Thus, this transcriptional state reflects an increased proliferative potential measured in vitro that is present in the normal esophagus and cancer.

## Discussion

In this study, we find that single epithelial cell clones have highly variable growth potential, with many clones that slow or halt their growth, but a rare subset that continues to grow exponentially. We find that this growth potential is heritable through cell division, suggesting that there are cell-intrinsic factors that govern growth potential. Furthermore, we find that the growth of clones can be modulated by altering cell density with increasing cell density leading to more homogeneous growth across clones. By analyzing transcriptional differences observed in clones of different sizes, we identify the specific cellular states that are differentially activated in the highly proliferative clones. Ultimately, we identify a similar highly proliferative cellular state within healthy human esophagus scRNA-seq in a subset of patients, with a notable increase in their prevalence within ESCC cases.

This study adds to the growing work on non-genetic cellular variability that is associated with distinct cell fates. The cell fate in this work is the proliferative capacity of each individual clone, which we measure by imaging individual clone sizes after a period of growth. We find that this growth fate is not encoded by genetic differences between cells and that both intrinsic and extrinsic factors can contribute to clonal growth fates. Using DNA barcoding technologies, we show that the cell states that mediate these growth behaviors are heritable through a limited number of cell divisions, similar to observations of memory in cancer drug resistance^26,29,38^, immune cells^46^, and differentiation^29,47,48^. While we describe the cell states associated with these phenotypic differences in epithelial progenitor cells, the mechanisms that encode the cellular state memory remain to be uncovered.

Our study uses an immortalized human esophagus epithelial cell line that recapitulates some but not all of the epithelial complexity seen in vivo. In mice, the esophageal epithelium is typically considered to be maintained by a single progenitor cell population that is thought to be functionally identical and equally capable of contributing to tissue maintenance^7,49,50^. Our in vitro system is different from such a model as we found that the individual progenitor clones have different growth potentials. In contrast, the human esophageal epithelium has two distinct progenitor cell populations^51–53^. These progenitor cells are thought to have different roles, with some primarily driving proliferation^51,54,55^, responding to environmental insults^56,57^, or maintaining genome stability^58,59^. Our in vitro model is likely more reflective of such complexity, and many of these processes could be contributing to the differences in clonal growth that we observe.

The relevance of our cell line model is supported by the identification of the same cell states in a rare subset of cells from the human esophageal epithelium. While we did not observe this signature in scRNA-seq from every patient sample, it is conceivable that this is due to the small size of the biopsied tissue. It is also possible that these cell states are associated with injury and regeneration, which would likely not be captured in every patient. Some of the patients had developed Barrett’s esophagus, but all biopsies from the healthy esophageal epithelium were taken far from the Barrett’s esophagus lesion. We then found this proliferative signature in a greater proportion of ESCC biopsies, suggesting its potential role in disease progression. Future studies will be needed to determine whether these cell states have functional consequences for driving growth in the normal and diseased esophagus.

In summary, we find clonal differences in the proliferative capacity of esophageal epithelial progenitors that are determined non-genetically. This study adds to a growing body of work across diverse phenotypes that had previously been explained exclusively by genetic mutations. In the case of the esophagus, clones can enter an expanding cell state that disrupts the balance between proliferation and differentiation. Future work is needed to determine whether these cell phenotypes and molecular states exist in epithelial cells beyond the esophagus.

## Methods

### Cell lines and culture

Human esophageal epithelial cells (EPC2-hTERT) were a gift from A. Muir (Department of Pediatrics at Children’s Hospital of Philadelphia, USA). We cultured EPC2-hTERT in Keratinocyte SFM (KSFM) supplemented with 2.5 µg of human recombinant epidermal growth factor (rEGF) and 25 mg of bovine pituitary extract (BPE) (Gibco, 17005042). Cells were passaged at 80% confluency using TrypLE Express Enzyme (Gibco, 12604013) and neutralized with equal amounts of 0.5mg/mL soybean trypsin inhibitor (STI; Gibco, 17075029). Human embryonic kidney cells (HEK293T) were cultured in 10% DMEM (90% DMEM high glucose with GlutaMAX (Gibco, 10569010), 10% FBS (GE, SH30396.03), 50 Units/mL penicillin, and 50µg/mL streptomycin (Gibco, 15070063)) and passaged with 0.05% trypsin-EDTA (Gibco, 25300054). All cells were incubated at 37°C and 5% CO_2_.

### Clonal isolation

We cultured EPC2-hTERT in KSFM supplemented with human rEGF and BPE. Cells were passaged two times before diluting to 0.5 cells per well in a 96-well plate. After 8 hours the cells adhered to the plate and each well was inspected and labeled for single cells. The medium was changed every 4 days and cells were grown for a total of 14 days. At this point, the clones had grown enough to become confluent in the well. A single confluent clone was then passaged and diluted to 0.5 cells per well and the same process was repeated. From observing the clones at multiple time points we could distinguish between expanding and committed clones.

### Small molecule inhibitors and recombinant proteins

PI3K inhibitor (GDC-0941, Cayman, 11600-10) was reconstituted at 10mM in DMSO for the stock solution. Final concentration for PI3K inhibitor treatment was 2µM. Human recombinant DKK1 (rhDkk, R&D Systems, 5439-DK) was reconstituted at 100µg/mL in 0.1% bovine serum albumin (BSA; Sigma-Aldrich, A7906). Final concentrations for PI3K inhibitor treatment were 200 ng/mL and 500 ng/mL.

### Immunofluorescence and phalloidin stain

EPC2-hTERT were washed twice with 1X Dulbecco’s phosphate buffered saline (DPBS; Corning, 21-031-CV) and fixed with 3.7% formaldehyde (Fisher, BP531-500) in PBS (Invitrogen, AM9625) for 10 min at room temperature. We then washed twice with PBS and permeabilized with 70% ethanol for at least 1 hour. Cells were blocked with 1% BSA in PBS for 1 hour at room temperature. Primary Ki-67 (SP6) rabbit monoclonal antibody (Sigma-Aldrich, SAB5500134) was diluted in 0.1% BSA in PBS (1:200 dilution). Primary staining solution was added and incubated overnight at 4°C. Secondary Alexa Fluor 647 goat anti rabbit IgG H+L (A21244) was diluted in 1% BSA in PBS (1:500 dilution) and combined with 7.5 µL stock solution of Phalloidin-Atto 488 (Sigma-Aldrich, 49409) for a total of 1.5mL 1% BSA in PBS. After two PBS washes, the secondary staining solution was added for 1 hour at room temperature in the dark. After two more PBS washes, we added 2x saline-sodium citrate (SSC) and stored at 4°C for imaging.

### Design gRNA plasmids

We designed CRISPR RNA (crRNA) in Benchling as a 23-nt sequence using variable protospacer adjacent motifs (PAMs) 5′-TTTV (TTTA, TTTC, or TTTG). 3 crRNA targeting functional domains were designed per gene. We used the AsCas12a crRNA expression vector (pRG212, Addgene_149722) to clone the guide RNA (gRNA). We digested and linearized the pRG212 vector with BsmBI (NEB, R0580), and the sense and anti-sense crRNA oligos were annealed and phosphorylated with T4 PNK (NEB, M0201). The linearized pRG212 vector and crRNA oligo were then ligated with T4 DNA Ligase (M0202).

### Generation and validation of Cas12 expressing cell line

We obtained the AsCas12a crRNA expression vector directly from Addgene (pRG232, Addgene_149723)^60^. The plasmid was packaged and transduced into HEK293T as described in the “Lentiviral packaging and transduction” section. Two days after transduction we passaged the cells to 2 10cm plates. The cells containing the Cas12-pRG232 insertion were selected with 0.5ug/mL puromycin (Takara, 631305) for three days. We determined the efficiency of the Cas12-pRG232 in our cell line by a competition-based proliferation assay. We generated two gRNAs linked to a GFP reporter; a positive control that targeted an essential gene (*PCNA*) and a non-targeting negative control (Rosa26)^60^. Cells were then transduced with either guide at 50% infection efficiency and the GFP-positive population was monitored using the BD Accuri C6 Plus Flow Cytometer every two days. We quantified editing efficiency by dividing the final GFP% timepoint by the initial GFP% timepoint per guide. We found that gRNA-Rosa26 stabilized around 98% and gRNA-*PCNA* drops to below 1%.

### Lentivirus transduction

To transfect HEK293T cells with the lentiviral plasmid, we first grew them to 90% confluence in a 10cm dish. We then prepared two tubes of transfection reagents: one containing 500µl of OPTI-MEM and 80 µl of 1 mg/mL PEI, and another containing 500µL of OPTI-MEM, 9 µg of psPAX2 plasmid, 5.5 µg of VSVG plasmid, and 8 µg of the barcode plasmid. After combining the contents of the two tubes, we incubated the mixture at room temperature for 20 minutes before adding it dropwise into the HEK293T plate. Following incubation of the cells at 37°C for 7 hours, we removed the medium and washed the plate once with DPBS. Next, we applied 6 mL of fresh medium and incubated the cells at 37°C for approximately 12 hours before repeating the medium collection process every 12 hours for a total of 72 hours. We filtered all the collected medium through a 0.2 µm filter and stored it at -80°C.

### Lentiviral transduction

Cells were transduced by creating a mixture consisting of polybrene (final concentration of 4 µg/mL), virus (determined by titration to be 20% infection), and cells at a concentration of 200,000 cells/mL. We then added 2 mL of this mixture to each well of a 6-well plate and subjected the plate to centrifugation at 600 RCF for 25 minutes. We incubated the cells with the virus at 37°C for 12 hours, after which we removed the virus-containing media and washed with DBPS. Subsequently, we added 2 mL of fresh medium to each well. The following day, we transferred each well to a 10 cm dish and allowed the cells to express the construct for 2-3 days.

### Fluorescence-Activated Cell Sorting (FACS)

We used TrypLE to detach the cells from the plate and STI to inhibit the TrypLE. The dissociated single-cell suspension was then washed once with 0.1% BSA and resuspended with 1% BSA and kept on ice. All cells that were sorted had a fluorescent tag that was detectable by the Beckman Coulter Moflo Astrios. We used a 100 µm nozzle and forward and side scatter to gate for single cells. We then sorted out the cells expressing the fluorescent tag, and WT cells were used as a negative control to determine gatting.

### Single-cell RNA sequencing

The library preparation was done in accordance with the manufacturer’s protocol (10x Genomics Chromium Next GEM Single Cell 3’ v3.1 Dual Index kit). For the esophageal epithelial cells (data shown in Fig. 3B,C), we sorted out 3,600 cells with lineage barcodes (GFP positive cells) and let them expand for 7.5 days. We then loaded the single-cell suspensions onto a 10x Chromium Controller to generate GEMs for each sample. We aimed to generate approximately 20,000 cells between two lanes while taking into account the estimated recovery of 50%. Finally, we sequenced all our libraries with paired-end sequencing on an Illumina NextSeq 550 using 10 cycles for both indices, 28 cycles for Read 1, and 43 cycles for Read 2.

### 10x Genomics sequencing data mapping and count matrix generation

The generation of FASTQ files was performed using CellRanger mkfastq v5.0.0, which processed raw Illumina base call files. The resulting FASTQ files were then aligned to the 10x reference GRCh38-2020-A using STARsolo v2.7.9a^61^. For downstream analyses, count matrices were generated utilizing the -soloFeatures GeneFull argument.

### scRNA-seq analysis and filtering

The count matrices for EPC2-hTERT were generated as mentioned above. scRNA-seq datasets of healthy human esophageal epithelium ^9,42–44^ and patients with ESCC ^44,45^ were previously published. Each dataset was individually filtered based on standard filtering before combining them using Seurat V4 ^62^. First, we excluded poorly sequenced cells with a low number of genes detected. Additionally, we removed cells that were likely to be doublets, characterized by a high number of genes. We then removed cells based on their mitochondrial percentage to eliminate low-quality or dying cells from our dataset. We also used a doublet finder (scDblFinder) to further filter doublets^63^. Each individual dataset was then normalized with “NormalizeData” and then all datasets were combined using “merge”. Finally we visualized the data with FindVariableFeatures, ScaleData and RunPCA. This was followed by RunUMAP (performing reductions with Harmony that significantly improve batch effects^64^), FindNeighbors, and FindClusters. Finally, we selected the expanding gene signature set by including *FOXM1, DKK1, MKI67*, and *TPX2* that are all a positive log_2_ fold-change differentially expressed genes between expanding and committed clones in our EPC2-hTERT model (Table S1).

### scMemorySeq: lineage barcode recovery from scRNA-seq and gDNA

Clonal dynamics were extracted using a high-complexity transcribed barcode library as previously described^26,37,38^, with the library generation protocol available at: https://www.protocols.io/view/barcode-plasmid-library-cloning-4hggt3w. Briefly, a CRISPR-Cas9 guide RNA plasmid (LRG2.1T) was used as a backbone for the DNA barcode. The plasmid was altered by replacing the U6 promoter and the single guide RNA scaffold with an EFS promoter and GFP. This was preceded by 100 semi-random nucleotides consisting of WSN repeats (W = A or T, S = G or C, N = any) that created the barcodes. The final sequence for the DNA barcode plasmid can be found here: https://benchling.com/s/seq-DAMUWPyU198hRSbpiecf.

The lineage barcodes are recovered from the excess full-length cDNA aliquot produced from the 10x library protocol. We PCR amplified the lineage barcodes with primers that were compatible with the NextSeq 500 and that would amplify the lineage barcode and the 10x cell barcode. The amplification was done by combining 100 ng of full-length cDNA per reaction, 0.5µM of each primer, and PCR master mix (NEB, M0543S). The amplification began with a 30-second denature step at 98°C, then 98°C for 10 seconds followed by 65°C for 2 minutes repeated 12 times, and a final extension step of 5 minutes at 65°C. We then performed SPRI bead (Beckman Coulter, B23317) size selection (using a 0.6X bead concentration) to extract the amplified barcodes (∼1.3kb). Finally, the lineage barcode library was sequenced with a NextSeq 500 with a Mid Output Kit v2.5 (150 cycles, Illumina, 20024904) by paired-end sequencing using 8 cycles on each index, 28 cycles on read 1 to read the 10x barcode and UMI, and 123 cycles on read 2 to sequence the lineage barcode.

The scRNA-seq and lineage barcode data was combined following the scMemorySeq custom pipeline that creates custom FASTQ files for the lineage barcodes that can be inputted in conjunction with the 10x Genomics Cell Ranger Feature Barcode pipeline. The custom python script is available through GitHub: https://github.com/SydShafferLab/BarcodeAnalysis.

Total genomic DNA was extracted from the samples using the QIAamp DNA Mini kit (Qiagen, 56304) following the manufacturer’s instructions. The target barcode region was amplified using PCR with primers targeting the region of interest that contained the Illumina adapter sequence, and index sequences. PCR amplification was carried out in a reaction mixture containing PCR Master Mix (NEB, M0543S), 0.5 μM of each primer, and 500 ng of template DNA. The amplification protocol consisted of an initial denaturation step at 98°C for 30 seconds, followed by 24 cycles of 95°C for 10 seconds and 65°C for 40 seconds, with a final extension at 65°C for 5 minutes. The PCR products were size excluded with SPRI beads (Beckman Coulter, B23317) to prepare the DNA library for sequencing using a double-sided clean-up. We then sequenced on an Illumina NextSeq 500 with a Mid Output kit (150 cycles, Illumina, 20024904) for single-end sequencing (151 cycles for read 1, and 8 cycles for each index).

### Determining expanding and committed clones for single-cell analysis

DNA barcodes were used to determine the expanding clones in the scRNA-seq data. We used the top 10% largest clones as our threshold for expanding colonies. We then used differential gene expression analysis between these newly labeled expanding cells and the remaining cells (Table S1). This highlighted a separate population with a unique gene expression state that we labeled as the committed clones. This annotation is supported by the DNA barcodes, since the smallest clones are captured by this group (Fig. 3E).

### Cell and colony counting

To evaluate cell counts post-fixation, we utilized Cellori, a nuclei counter which can be accessed at https://github.com/zjniu/Cellori/tree/main and DeepCellHelper (https://github.com/SydShafferLab/DeepCellHelper). To identify individual clones form the output of Cellori and DeepCellHelper, we developed a custom graphical user interface, ColonySelector, which can be found at https://github.com/SydShafferLab/ColonySelector. By circling individual clones in each well using ColonySelector, we generated a comprehensive record associating each nucleus with its respective colony. Through the integration of these methodologies, we achieved a precise and efficient analysis of clonal counts.

## Data and code availability

All code and data used will be made available upon publication.

## Acknowledgments

We thank all members of the Shaffer lab for feedback on experiments and the manuscript, A. Singh for thoughtful discussions and ideas, G. Harmage for help on developing image processing and barcoding pipelines, and A. Muir for providing cell lines. S.M.S. acknowledges support from the NIH Director’s Early Independence Award DP5OD028144, a donation to the Institute for Regenerative Medicine at the University of Pennsylvania from Larry and Mickey Magid, and the Institute for Translational Medicine and Therapeutics of the Perelman School of Medicine at the University of Pennsylvania (NIH NCATS UL1TR001878). R.A.R.H. acknowledges support from the NSF Graduate Research Fellowship (DGE-1845298).

## Author contributions

Conceptualization, R.A.R.H., R.A.G. and S.M.S.; Methodology, R.A.R.H., R.A.G. and S.M.S.; Investigation, R.A.R.H. and R.A.G.; Software, R.A.R.H. and R.A.G.; Formal Analysis, R.A.R.H.; Data Curation, R.A.R.H. and R.A.G.; Writing – Original Draft, R.A.R.H. and S.M.S.; Writing – Review and Editing, R.A.R.H., R.A.G and S.M.S.; Visualization, R.A.R.H.; Funding Acquisition, S.M.S.; Supervision, S.M.S.

