## Supplementary Figures for "Non-genetic differences underlie variability in proliferation among esophageal epithelial clones"

**R. A. Reyes Hueros et al.**

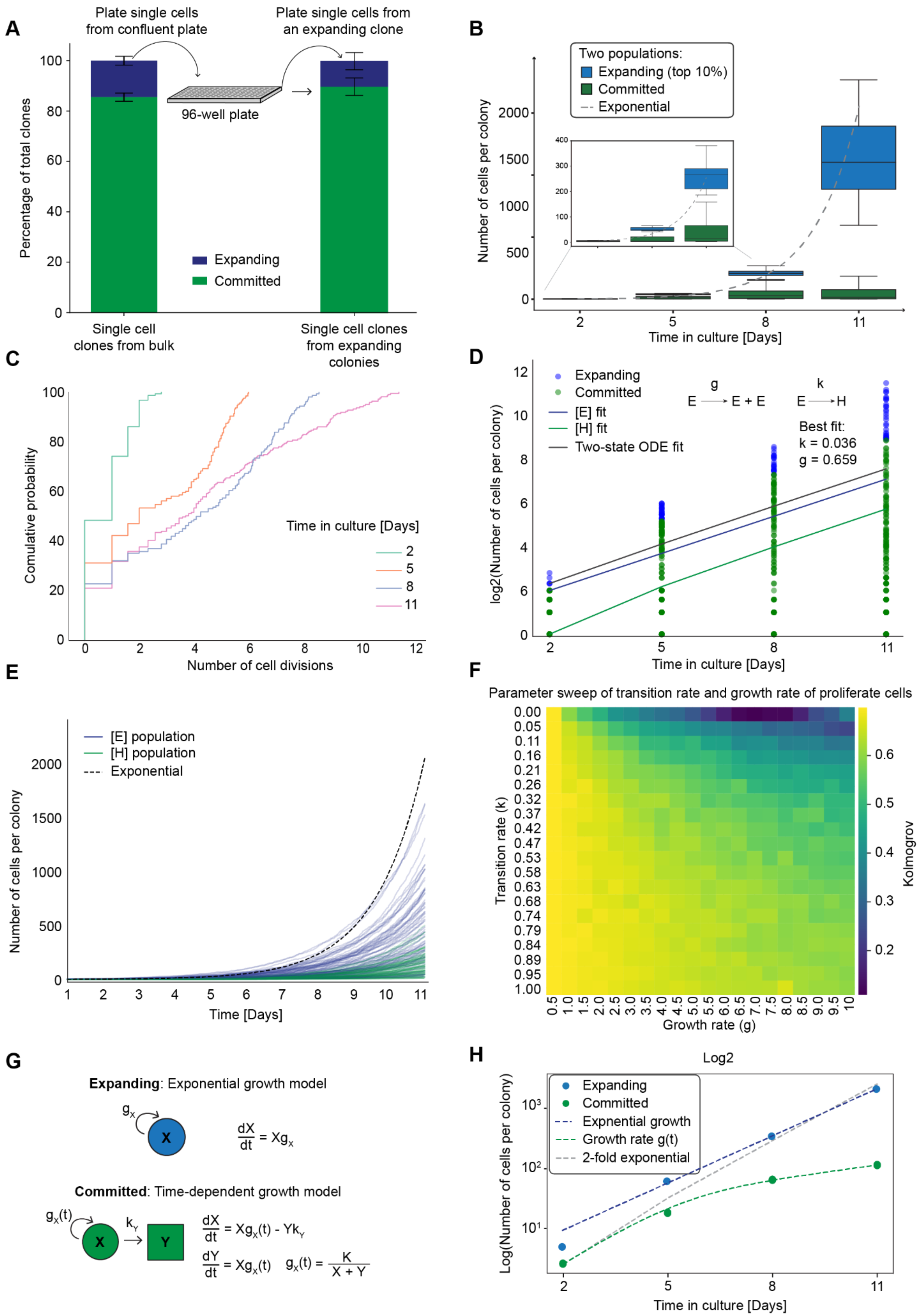
A
Plate single cells from confluent plate
Plate single cells from an expanding clone
96-well plate
Percentage of total clones
Expanding
Committed
Single cell clones from bulk
Single cell clones from expanding colonies
B
Two populations:
Expanding (top 10%)
Committed
Exponential
Number of cells per colony
Time in culture [Days]
C
Comulative probability
Time in culture [Days]
Number of cell divisions
D
Expanding
Committed
[E] fit
[H] fit
Two-state ODE fit
E → E + E
E → H
Best fit:
k = 0.036
g = 0.659
log2(Number of cells per colony)
Time in culture [Days]
E
[E] population
[H] population
Exponential
Number of cells per colony
Time [Days]
F
Parameter sweep of transition rate and growth rate of proliferate cells
Transition rate (k)
Growth rate (g)
Kolmogrov
G
Expanding: Exponential growth model
Committed: Time-dependent growth model
H
Log2
Expanding
Committed
Expnential growth
Growth rate g(t)
2-fold exponential
Log(Number of cells per colony)
Time in culture [Days]

**Fig. S1. Mathematical models for epithelial cell growth. (A**) Single EPC2-hTERT cells were initially plated in individual wells of a 96-well plate and their growth monitored over a 14-day period. The left bar plot shows the majority of clones slowed or ceased their growth, while approximately 10% of clones exhibited exponential growth. To investigate whether this observed clonal heterogeneity was due to genetically distinct subclones, the most proliferative clones were isolated and subjected to a second round of single cell plating. The right bar plot shows subclones derived from highly proliferative parent clones match the initial population distribution, negating the idea of genetic differences driving growth disparities and emphasizing the inherent variability in EPC clonal growth capacities. (**B**) Bar plot showing cell divisions per clone over time, with expanding (blue) and committed (green) cells. A solid gray line highlights exponential growth for comparison. Expanding cells represent those that are actively dividing and contributing to the overall growth of the clone (10%), whereas Committed cells have slowed their proliferation. By comparing the number of cell divisions for these two populations across different days in culture, the plot provides insights into the dynamics of cell proliferation and differentiation over time. (**C**) Cumulative distribution function plot of the number of cell divisions per clone separated by days in culture. A subset of clones continue to proliferate as described by the growth models for expanding cells. (**D**) An optimization technique (minimize the sum of squared differences) was used to fit the model parameters to the data. The simple two state ODE that we are fitting is depicted in the top right and the best fit parameters for the growth and transition rate are below. (**E**) A Gillespie stochastic simulation was used to capture the inherently stochastic nature of biological processes described by the two-state model that lead to a broad range of clone sizes. (**F**) We compared the simulated and experimental growth distributions using the Kolmogorov-Smirnov test. These tests provided a quantitative measure of the similarity between the simulated and experimental distributions, allowing us to assess the goodness of fit of the models and parameter sets. The results of these tests consistently supported the findings that a more complex model might be necessary to fully capture the growth behavior of the cells. (**G,H**) The data was then split as in expanding and committed and modeled separately. The left panel displays growth models for expanding and committed clones, with expanding clones fitted using an exponential growth model and committed clones requiring a time-dependent model for accurate data representation. Growth models are provided within the figure. The right panel shows the data fitted by their respective growth rate models, illustrating the decaying growth of committed clones and the constant growth rate of expanding clones.

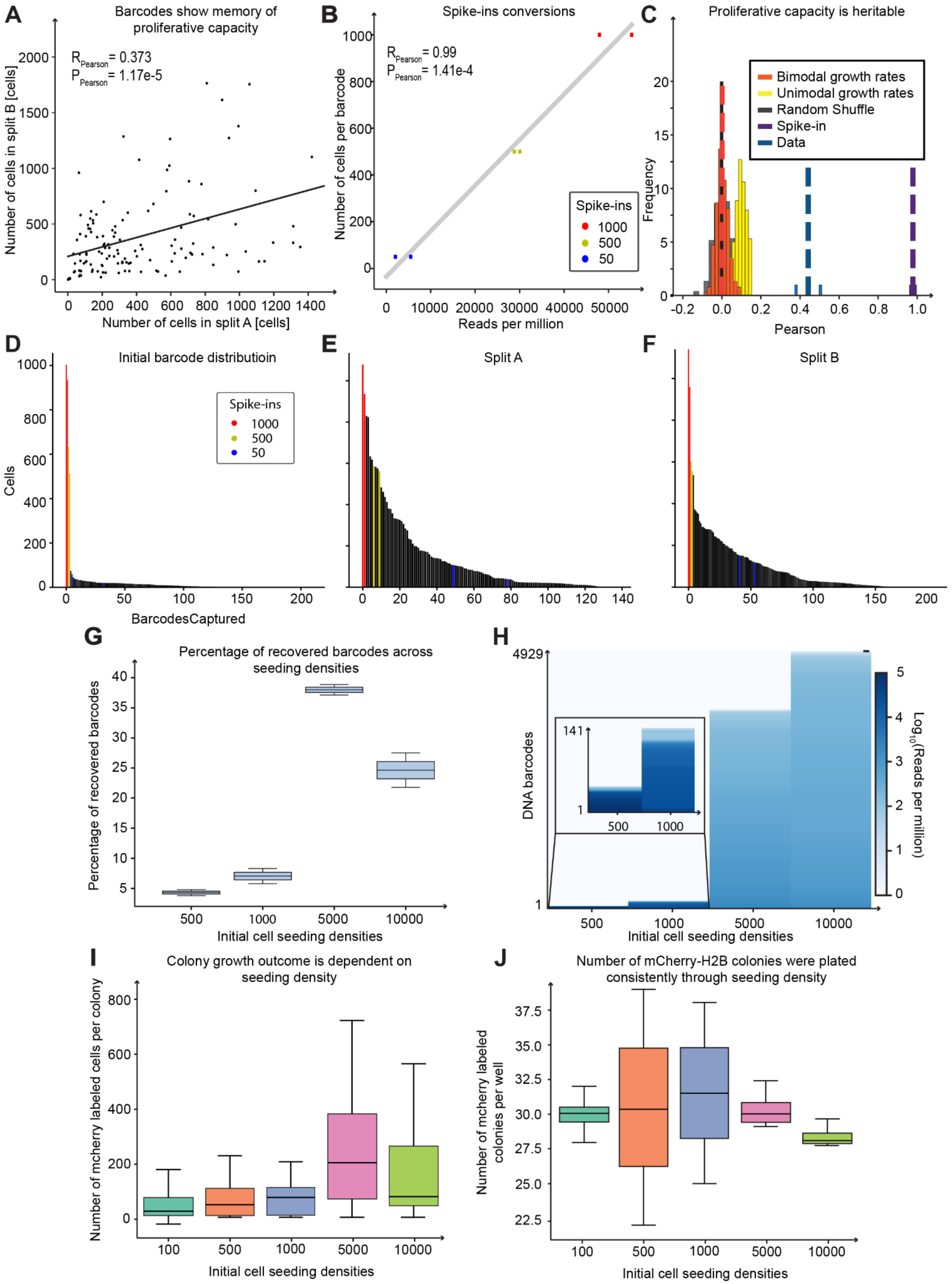

A
Barcodes show memory of proliferative capacity
$R_{Pearson}$ = 0.373
$P_{Pearson}$ = 1.17e-5
Number of cells in split B [cells]
Number of cells in split A [cells]
B
Spike-ins conversions
$R_{Pearson}$ = 0.99
$P_{Pearson}$ = 1.41e-4
Number of cells per barcode
Reads per million
Spike-ins
1000
500
50
C
Proliferative capacity is heritable
Bimodal growth rates
Unimodal growth rates
Random Shuffle
Spike-in
Data
Frequency
Pearson
D
Initial barcode distributioin
Spike-ins
1000
500
50
Cells
BarcodesCaptured
E
Split A
F
Split B
G
Percentage of recovered barcodes across seeding densities
Percentage of recovered barcodes
Initial cell seeding densities
H
DNA barcodes
4929
141
$Log_{10}$(Reads per million)
Initial cell seeding densities
I
Colony growth outcome is dependent on seeding density
Number of mcherry labeled cells per colony
Initial cell seeding densities
J
Number of mCherry-H2B colonies were plated consistently through seeding density
Number of mcherry labeled colonies per well
Initial cell seeding densities

**Fig. S2. Evidence that proliferative potential is heritable through cell division.** (**A**) A scatter plot comparing clonal cells in split A and split B revealed a Pearson correlation of 0.373 and a highly significant p-value of 1.17e-5. This relationship, observed in the second replicate (2/2), suggests a true proliferative correlation between the clonal cells. (**B**) Scatter plot of spike-ins used in split barcode experiment. Spike-ins are an external reference for converting transcript counts to cell numbers and were added to each sample during library preparation (see methods). Spike-ins consist of known barcode sequences, where a specific quantity of them are added to normalize transcripts across samples. The relative abundance of transcripts was quantified by comparing transcript counts to spike-in counts, enabling the conversion of transcript counts to cell numbers. (**C**) Histograms of Pearson correlations for simulated data, split barcode experiments, and spike-ins. Simulated data includes three scenarios: bimodal growth, unimodal growth, and stochastic growth, each based on distinct growth rate probability distributions. Bimodal growth uses a bimodal fit from day 8 data, unimodal growth employs a Gaussian curve, and stochastic growth assigns equal probability to all experimental growth rates. The null hypothesis assumes randomly chosen growth rates for sibling cells, eliminating memory of their prior proliferative state. Both replicates, shown in blue, fall outside the simulation ranges, indicating memory of epithelial cell proliferative capacity. Spike-ins serve as highly correlated positive controls by design. **(D-F**) Bar plots of rank-ordered barcodes, representing initial barcode distribution and barcode distribution after 8 days of growth. Reads Per Million were converted to total cell counts using spike-ins. Spike-ins are displayed in red, yellow, and blue, corresponding to 1000, 500, and 50 cells, respectively. (**G)** Description for box plot with seeding density: We plated barcoded cells at different densities and allowed them to grow for 8 days. We then collected the barcodes and quantified the size of the clones based on the number of reads of their barcodes. We found the largest clone sizes at 5000 cells per well. Box plots show the percentage of clones detected. In these experiments, growth correlates with our ability to capture the barcode by sequencing, and bigger clones are more likely to be captured with sequencing. (**H**) Rank-ordered barcodes grouped by density. The heat bar of log(Reads per Million) shows that the read depth of the barcodes is equal throughout distinct densities. (**I**) Box plot of data from Fig 2D. Using mCherry labeled cells and quantifying clone growth as described in Fig. 2C, we further confirmed the results of Fig. S2G. The box plot shows the number of cells in each clone and confirms the largest clone size at 5000 cells per well. (**J**) Box plot showing accurate plating of the mCherry clones within the field of unlabeled cells. The same number of mCherry clones is present in each well.

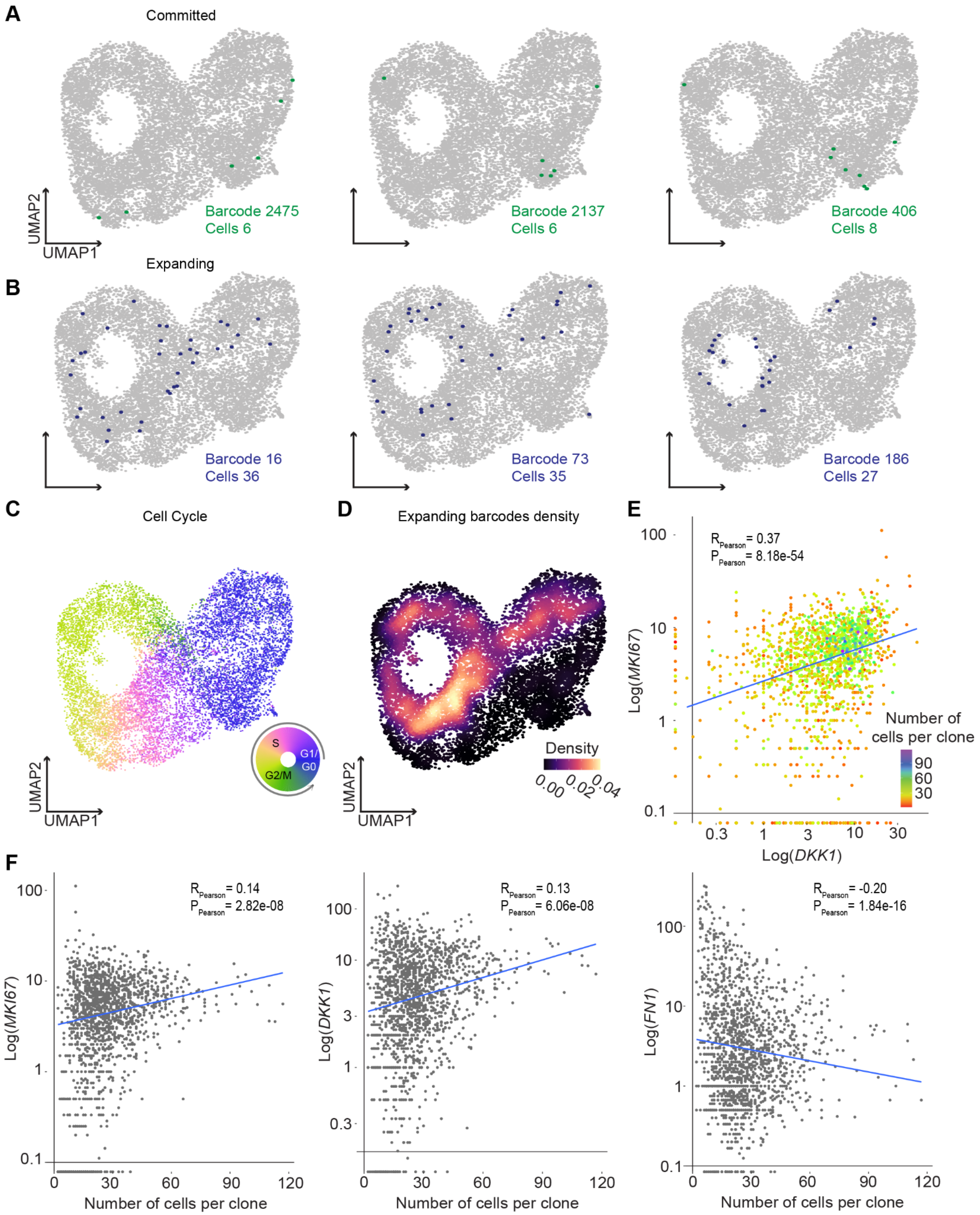

A
Committed
Barcode 2475
Cells 6
Barcode 2137
Cells 6
Barcode 406
Cells 8
UMAP2
UMAP1
B
Expanding
Barcode 16
Cells 36
Barcode 73
Cells 35
Barcode 186
Cells 27
C
Cell Cycle
S
G1/G0
G2/M
D
Expanding barcodes density
Density
0.00
0.02
0.04
E
R Pearson = 0.37
P Pearson = 8.18e-54
Log(MKI67)
Log(DKK1)
Number of cells per clone
90
60
30
F
R Pearson = 0.14
P Pearson = 2.82e-08
Log(MKI67)
Number of cells per clone
R Pearson = 0.13
P Pearson = 6.06e-08
Log(DKK1)
R Pearson = -0.20
P Pearson = 1.84e-16
Log(FN1)

**Fig. S3. Molecular characterization of expanding and committed clones. (A**) UMAP plots with examples of lineages from committed clones. (**B**) UMAP plots with examples of lineages from expanding clones. (**C**) UMAP embeddings, with cells colored based on their cell-cycle position using a circular color scale. Discrete stage labels are placed in approximate positions on the circular legend. (**D)** UMAP embeddings are overlaid with kernel density estimations showing locations of cells with an expanding signature. (**E**) Scatter plot of *MKI67* and *DKK1* log-scaled with a heat bar representing the number of cells per clone. *MKI67* and *DKK1* have a Pearson correlation of 0.37 and the top right quadrant has the largest clones. (**F**) Scatter plot of number of cells per clone against *MKI67*, *DKK1* and *FN1* log-scaled.

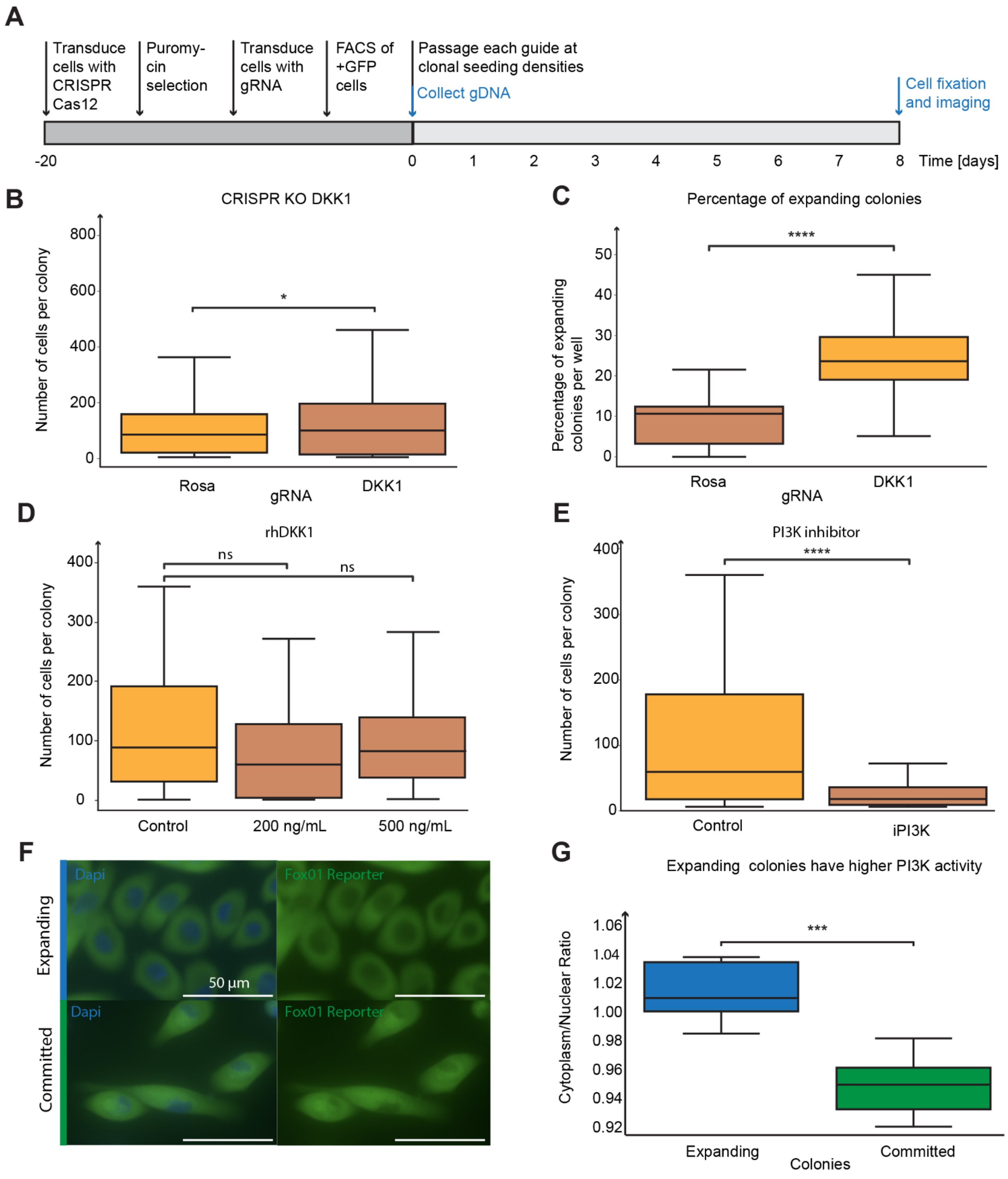

A
Transduce cells with CRISPR Cas12
Puromy-cin selection
Transduce cells with gRNA
FACS of +GFP cells
Passage each guide at clonal seeding densities
Collect gDNA
Cell fixation and imaging
-20
0
1
2
3
4
5
6
7
8
Time [days]
B
CRISPR KO DKK1
Number of cells per colony
800
600
400
200
0
*
Rosa
gRNA
DKK1
C
Percentage of expanding colonies
Percentage of expanding colonies per well
50
40
30
20
10
0
****
Rosa
gRNA
DKK1
D
rhDKK1
Number of cells per colony
400
300
200
100
0
ns
ns
Control
200 ng/mL
500 ng/mL
E
PI3K inhibitor
Number of cells per colony
400
300
200
100
0
****
Control
iPI3K
F
Expanding
Committed
Dapi
Fox01 Reporter
50 µm
G
Expanding colonies have higher PI3K activity
Cytoplasm/Nuclear Ratio
1.06
1.04
1.02
1.00
0.98
0.96
0.94
0.92
***
Expanding
Colonies
Committed

**Fig. S4. WNT and PI3K maintain the proliferative capacity of expanding clones. (A)** Outline of experimental design for the CRISPR-Cas12a targeted knockout (KO) approach is outlined, which is employed to investigate the role of DKK1 in clone expansion. (**B**) Box plot of percentage of expanding clones per well split by gRNA targeting Rosa26 and DKK1. (**C**) Box plot of percentage of expanding clones per well split by gRNA targeting Rosa26 and DKK1. DKK1 KO has a significant increase in expanding clones. (**D**) Box plot of the number of expanding clones between different concentrations of human recombinant DKK1; there is no significant effect on clonal growth. (**E**) The box plot reveals the number of cells per clone after a 7-day treatment with PI3K inhibitor (iPI3K). (**F,G**) Representative images of PI3K reporter activity in both expanding and committed clones are displayed on the left, offering a visual representation of PI3K activity in the two different cell populations. The PI3K reporter, FOXO1, offers higher specificity and an increased dynamic range for detecting PI3K pathway activity. Furthermore, the increased dynamic range of the FOXO1 reporter enables the detection of subtle changes in PI3K activity, allowing for a more comprehensive understanding of the pathway's role in various cellular processes. On the right, a box plot quantifies the PI3K reporter signal as the ratio of cytoplasmic to nuclear fluorescence, with a highly significant p-value of 2.37e-3 observed between expanding and committed clones. This suggests a distinct difference in PI3K signaling between the two cell populations, further emphasizing the roles of PI3K in maintaining the proliferative capacity of expanding clones.

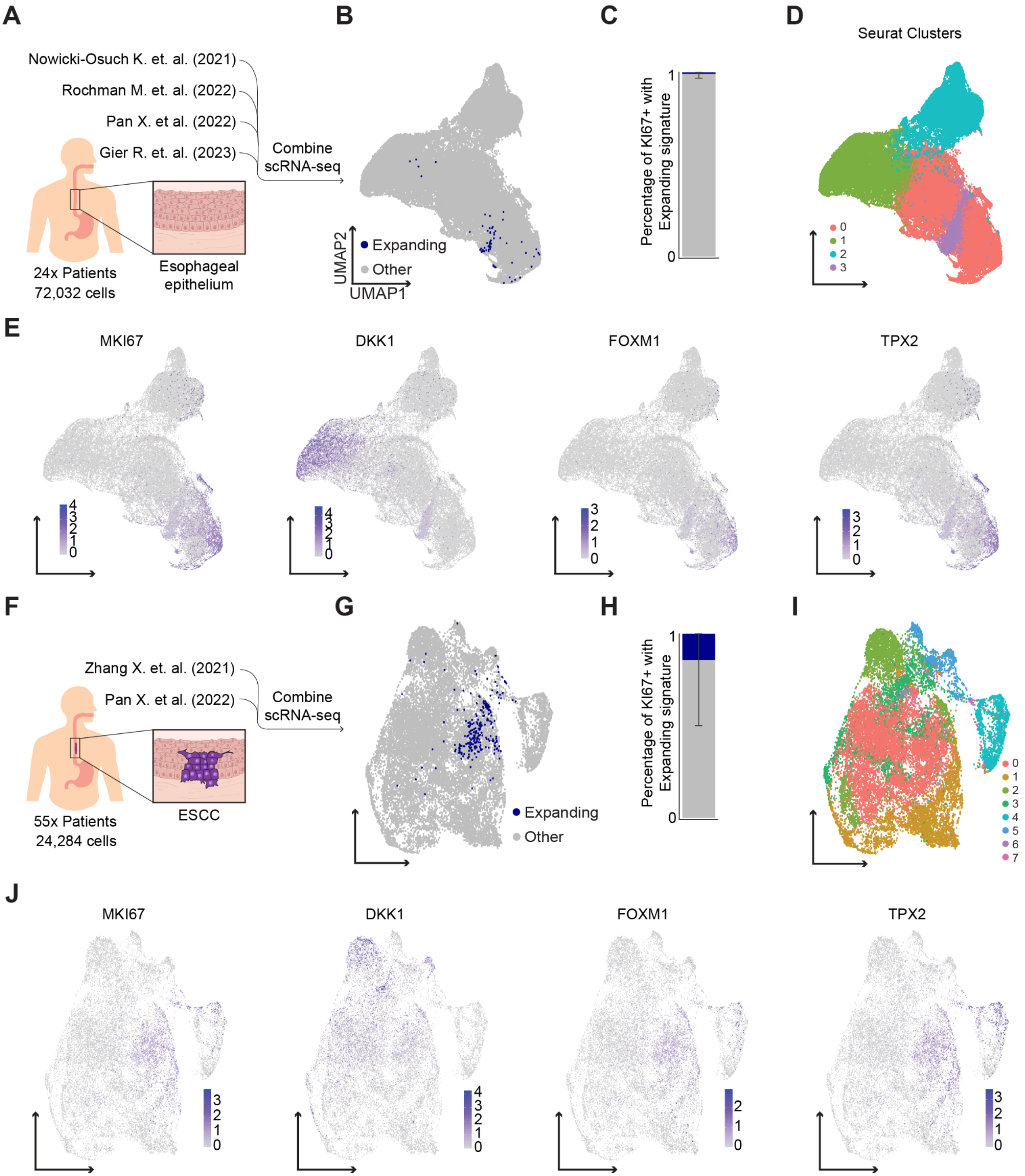

A
Nowicki-Osuch K. et. al. (2021)
Rochman M. et. al. (2022)
Pan X. et al. (2022)
Gier R. et. al. (2023)
Combine scRNA-seq
24x Patients
72,032 cells
Esophageal epithelium
B
Expanding
Other
UMAP2
UMAP1
C
Percentage of KI67+ with Expanding signature
D
Seurat Clusters
E
MKI67
DKK1
FOXM1
TPX2
F
Zhang X. et. al. (2021)
Pan X. et al. (2022)
Combine scRNA-seq
55x Patients
24,284 cells
ESCC
G
Expanding
Other
H
Percentage of KI67+ with Expanding signature
I
J
MKI67
DKK1
FOXM1
TPX2

**Fig. S5. Additional characterization of the expanding signature.** (**A**) A schematic of the integration of numerous published scRNA-seq datasets for healthy esophageal epithelium comprising 26 patients and 62,216 cells. (**B**) UMAP plot with cells expressing the expanding signature labeled in blue. (**C**) Bar plot showing the percentage of *MKI67* positive cells expressing the expanding signature. (**D**) UMAP plot of Seurat clusters. The clusters represent tissue layers: (0) basal layer, (1) superficial layer, (2) suprabasal layer, and (3) the cells that most closely match the expanding clone signature. (**E**) UMAP plots of the genes that make up the expanding cell signature. (**F**) A schematic of the integration of numerous public scRNA-seq datasets for ESCC with 65 patients and 19,289 cells. (**G**) UMAP plot with cells expressing the expanding signature labeled in blue. (**H**) Bar plot showing the percentage of *MKI67* positive cells expressing the expanding signature. (**I**) UMAP plot of Seurat clusters. (**J**) UMAP plots of the genes that make up the expanding cell signature.
